# Precision not prediction: Body-ownership illusion as a consequence of online precision adaptation under Bayesian inference

**DOI:** 10.1101/2024.09.04.611162

**Authors:** Filip Novický, Ajith Anil Meera, Fleur Zeldenrust, Pablo Lanillos

## Abstract

Humans can experience body ownership of new (external) body parts, for instance, via visuotactile stimulation. While there are models that capture the influence of such body illusions in body localization and recalibration, the computational mechanism that drives the experience of body ownership of external limbs is still not well understood and under discussion. Here, we describe a mathematical model of the dynamics of this phenomenon via uncertainty minimization. Using the Rubber Hand Illusion (RHI) as a proxy, we show that to properly estimate one’s arm position, an agent needs to infer the least uncertain world model that explains the observed reality through online adaptation of the signals’ relevance, i.e., its precision parameters (the inverse variance of the prediction error signal). Our computational model describes that the illusion is triggered when the sensory precision estimate quickly adapts to account for the increase of sensory noise during the physical stimulation of the rubber hand due to the occlusion of the real hand. This adaptation produces a change in the uncertainty of the body position estimates, yielding a switch of the perceived reality: the “rubber hand is the agent’s hand” becomes the most plausible model (i.e., it has the least posterior uncertainty). Overall, our theoretical account, along with the numerical simulations provided, suggests that while the perceptual drifts in body localization may be driven by prediction error minimization, body-ownership illusions may be a consequence of estimating the signals’ precision, i.e., the uncertainty associated with the prediction error.

## 1. Introduction

Humans navigate the environment and interact with it through their bodies. This happens with a precise balance between internal representations of our body learned over our lifetime [61] and sensorimotor-related processes, such as sensory signals arriving from the periphery caused by the localization of our body in space [17, 75]. Remarkably, these processes are “flexible”, probably to handle uncertainty and conflicts in the sensory input and to adapt to bodily changes and changes in the external world. Strong experimental evidence reveals that body perception can be easily manipulated [16, 44, 34]. Body-ownership, i.e., “perceptual experience of body or body part as one’s own” [17], has been associated with information integration in the ventral premotor cortex [19], right posterior insula, right frontal operculum as well as posterior parietal cortex junction [10, 31, 47]. However, the exact computational mechanisms behind and its intricate neural underpinnings are still not well understood. By developing computational models that replicate perceptual and motor effects of humans when subjected to multisensory conflicts, such as body illusions [18, 46], some inner characteristics of these hidden mechanisms may be revealed [44, 35]. In this sense, one of the most intriguing scientific questions is how we can experience the embodiment of external objects as a part of our own body.

To study this, we focused on the mechanism behind the body ownership illusion and its temporal dynamics [25]. While the research literature already provides computational accounts to explain body mislocalization as an effect of body illusions [44] and its dynamics [35, 46], there are no mathematical accounts that properly describe and explain the dynamics of body-ownership illusions. This is going beyond static Bayesian Causal Inference (BCI) [71, 9] and showing the temporal dynamics of the mechanism behind how an external object becomes part of your body during the illusion. Here, we provide a novel mathematical account that explains the effect of body-ownership illusions as an indirect consequence of uncertainty minimization. In particular, our model describes that perceptual drifts in body localization are driven by prediction error minimization, and body-ownership illusions are a consequence of online estimation of the signals’ precision. Our model shows that the illusion is triggered when the sensory precision estimate quickly adapts to account for the increase of sensory noise during the physical stimulation of the fake body part due to the occlusion of the real body part. This adaptation produces a change in the uncertainty of the body localization estimates, yielding a change in the perceived reality: the ‘rubber hand is the agent’s hand’ becomes the most plausible model—it is the hypothesis or model that has the least posterior uncertainty. Importantly, our model aligns with previous conceptual accounts of causal inference [71, 35] of body perception and the recent findings regarding the relation between the emergence of the illusion and visual noise [10, 9].

To introduce the proposed model, we first summarize previous research on limb illusions and related computational models that are the backdrop of our body ownership model based on precision.

### 1.1. Previous research on limb illusions and related computational models

There is an extensive body of literature on limb-body illusions and manipulations—see [44] and [18] for a review. Here, we briefly summarize the basic findings that point out two well-known phenomena: *i*) body-ownership of external/fake limbs and *ii*) limb mislocalization (or perceptual drift).

Evidence of the body-ownership of external/fake limbs shows that the majority of tested subjects—not all of them—perceive a fake limb as their own after a short period of a specific type of visuotactile stimulation [6, 44]. The most well-studied body illusion, the Rubber Hand Illusion (RHI) [6], can be seen as a phenomenon where people hold false beliefs about the causes of tactile stimulation. Specifically, the participant sees a rubber hand being stroked (e.g., with a brush) at the same time as the real hand, which is occluded. In less than one minute, the participant has the subjective experience of owning the fake hand as part of the body [6]. This illusion comes with appearance and positional constraints, and its intensity varies depending on the subject [45]. The illusion can be measured using subjective questionnaires. Objective measures, such as skin conductance, were introduced to evaluate the body’s reaction to the process of the fake limb embodiment [5, 33, 21] and confirmed that external hands also produce physical responses.

The second well-known phenomenon is perceptual drift, which can be seen as a mislocalization phenomenon due to sensory integration [74]. Participants localize their own limb slightly drifted towards the fake limb [75, 32]. By measuring the perceived location of the hand before and after the illusion is induced, researchers have suggested that the closer participants estimate their real hand to the position of the rubber hand, the stronger the illusion is [74]. While studies used this measure to elucidate the effects of the ownership effect [6, 14], other studies highlighted that the correlation between the perceived ownership and the proprioceptive drift does not necessarily emerge [37, 38, 67, 68, 1], questioning the use of perceptual drift to investigate the feeling of body ownership, and thus suggesting that perceptual drift and body-ownership illusion are a consequence of two interrelated but different processes.

Computational models of body perception should be able to account (at least) for the two well-known phenomena mentioned above. There are already candidate computational models for the mislocalization effect based on the optimal integration of the multisensory signals [71, 35, 46, 69, 49]. The rationale is that the brain optimally integrates tactile, visual and proprioceptive cues to make them fit into an internal model of the body (learned through experience). In the presence of multisensory conflicts induced by the new visuotactile input, the part of the internal model that is used to explain body localization and that is adaptive forces the system to incorrectly estimate the limb in the space. The models differ in the mathematical framework used to integrate the signals [44]: optimal signal integration [20], Bayesian Causal Inference [71], Free Energy Minimization [35, 46], and synaptic plasticity (e.g., spike-timing-dependent plasticity) [76].

The computational mechanism behind the body ownership illusion is more elusive than the mechanism behind the mislocalization effect. One of the most influential proposed mechanisms, from a conceptual point of view, is that the brain, during the body illusion experiment, decides whether the information comes from the fake limb or the own limb [71, 44]. This means that the subject decides between two models of the perceived reality. This may explain why the increase of the number of presented cues (visuotactile synchronization, coherent poses, active movement) that support the fake limb model results in a stronger illusion. Thus, according to this computational model, causal inference plays a strong role in the body illusion. Still, the computational process and its dynamics behind the perceptual experience where the participant starts feeling a fake limb as its own remains unknown. Bayesian Causal Inference, as presented in [71], describes body ownership as a static probability computation, where no explanation is given on the inner functional mechanism producing the dynamics of the perceptual drifts nor the illusion [25]. Importantly, in recent experiments, the notion of uncertainty as a driving mechanism due to (visual) noise induction [10, 9] is gaining strength. It aligns with a hypothesis that body ownership works as a perceptual inference process, where uncertainty monitoring modulates the experience.[46]. For a more comprehensive summary of computational models of body illusions, see the Appendix 5.1.

### 1.2. A unified body-ownership model based on precision

The models described in the previous section vary significantly in their approaches to investigating specific phenomena observed in limb embodiment illusions. However, none have yet fully explained the mechanism and dynamics behind the ownership illusion and its relation to multisensory integration and localization drifts. We address this gap by presenting a novel mathematical description of the mechanism that may trigger the feeling of owning a fake limb. Our model builds upon previous seminal Bayesian modeling accounts (i.e., Predictive Processing [46] and Bayesian Causal Inference [71]), but focuses on the role of the precision (inverse variance) adaptation during body illusions. In Bayesian models, the sensory precision parameter determines the weight of prediction errors arising from different sensory modalities. These prediction errors are, in turn, used in attention and uncertainty processing in the brain [58, 59]. Prior work suggests that precision optimization may be critical for resolving conflicts between competing models of sensory input [10]. However, existing computational accounts have not fully explored how dynamic precision adaptation could explain the temporal evolution of body ownership illusions. Understanding this process could reveal how the neuronal processes actively adjust the relative influence of different sensory signals to resolve uncertainty when faced with conflicting bodily information.

Our hypothesis posits that body illusions are a consequence of the brain aiming to minimize uncertainty by (online) adapting to the precision of prediction errors. Neurobiologically, this precision parameter can be related to the postsynaptic gain, which influences the encoding of prediction errors [54, 24, 3, 28, 4]. Interestingly, aberrant precision has been implicated in the causes of hallucinations [7, 40, 23, 39, 3], and these precisions are thought to be modulated via NMDA receptors and neuromodulators such as dopamine, acetylcholine, serotonin, and noradrenaline [3, 58, 57, 56, 55]. Essentially, as precision reflects the uncertainty of the prediction error, it may also function as a model of the experimental results observed when introducing visual noise in a body illusion experiment, where sensory uncertainty was found to be essential for the ownership illusion [10].

We propose a computational model of body ownership illusion – evaluated in the RHI paradigm – that uses precision and its adaptation to sensory noise levels to explain the perceptual switch of feeling a fake hand as part of the body. The model schematic, the paradigm and used variables are described in Figure 1A,B and C, respectively. Similarly to the Bayesian Causal Inference approach [71], the brain decides between two different hypotheses or models *M* of reality. However, our approach will base its decision on each model’s posterior uncertainty. Model *M*_1_ explains the observed data as “my hand is the real hand”, while *M*_2_ explains the data as “my hand is the rubber hand”. The induced noise from covering the real hand prompts the system to adapt. This adaptation makes the posterior estimates more uncertain. Consequently, hypothesis/model selection switches to the most probable one, which is to own the fake limb, thus producing the illusion.

**Fig 1:**
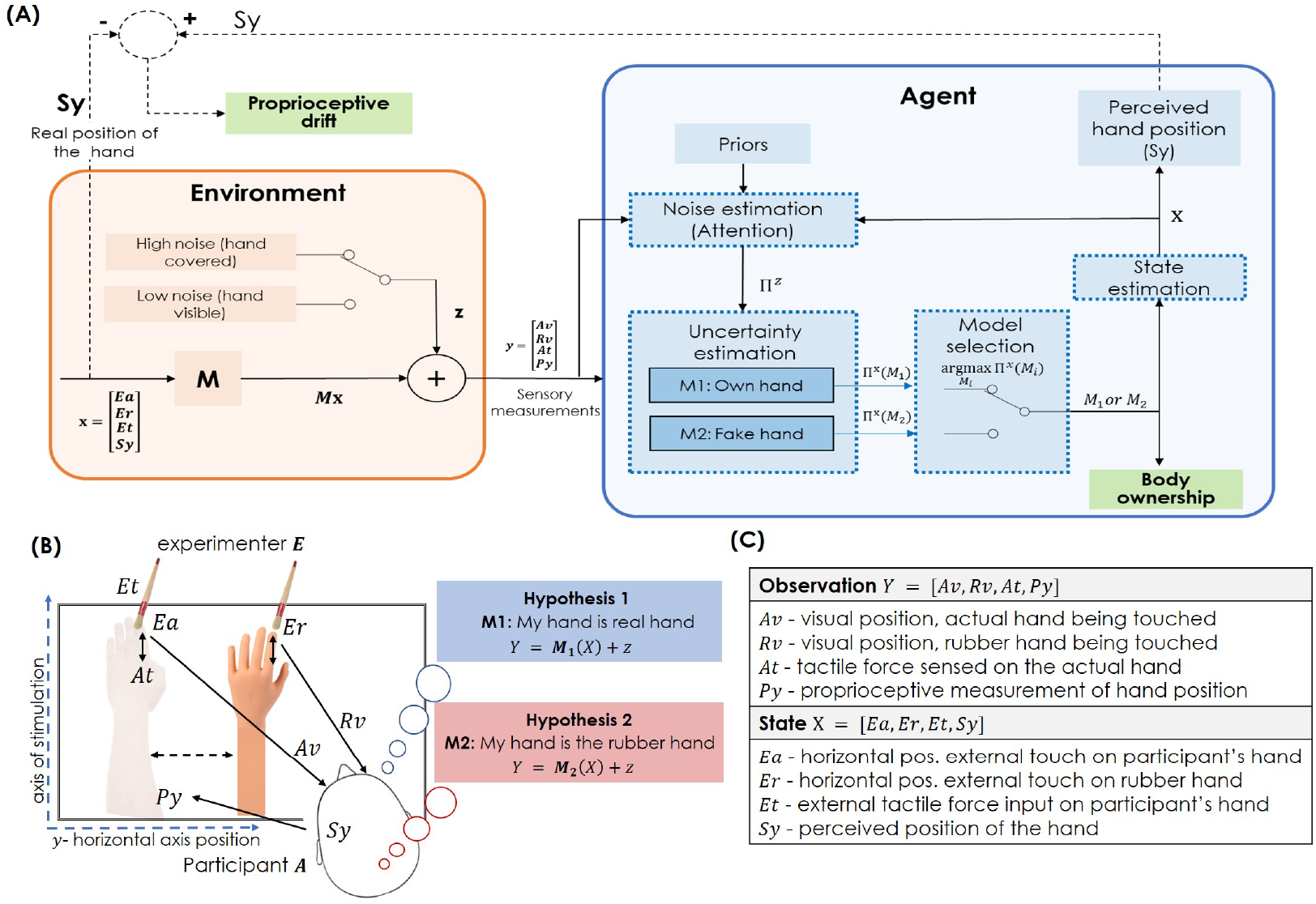
Maximum posterior precision body-ownership model. (A) Our proposed model consists of two main blocks: the environment and the agent. The state of the environment *x* produces the output *y* that is measured by the agent as sensory measurements, along with the observation noise *z*. The visual noise is high when the real hand is covered, and low when the real hand is visible. The agent estimates the noise via attentional mechanisms, i.e., precision priors, which leads to estimating the uncertainty of the *M*_1_ and *M*_2_ models. Through model selection, the agent selects the best suitable model, producing the body-ownership illusion. Using the selected model, the state estimation is performed, resulting in the state estimate *X* that represents *x*. The proprioceptive drift is the difference between the perceived hand position *S y* and the real position of the hand **Sy** (that the agent does not have access to). Note that all variables in **bold** belong to the generative process (such as **Sy**), while non-bold variables belong to the generative model (such as *S y*). (B) Experiment formalization. The brain is testing two competing models (i.e., hypotheses): *M*_1_ stands for believing that “my hand is the real hand”, while *M*_2_ indicates that “my hand is the rubber hand”. These two internal models maps (as a generative model) the states *X* into the observations *Y*. (C) Notation used for the observations (sensory measurements) and the states.

Our computational model suggests that: *i*) body mislocalizations (e.g., proprioceptive drift) result from body state estimation, where prediction errors are minimized, and *ii*) body ownership illusions, i.e., perception of a fake limb/hand as one’s own, result from uncertainty minimization, where precision adaptation to overcome induced noise triggers a model selection switch.

## 2. Methods

We introduce a mathematical model for the body-ownership illusion, which is drawn from the Bayesian brain hypothesis [15] and grounded on the free energy principle (FEP) [29]. This principle proposes that the brain generates cognition and behaviour following a Bayesian optimization strategy. It postulates that living beings show resilience over time, which is achieved via minimizing entropy (the second law of thermodynamics) to preserve their equilibrium state (i.e., to self-organize) [27, 60]. To do this, living systems embed a generative internal model that explains the world (i.e., perception) and adjusts to the environment (by changing their belief or exerting actions) via minimizing the free energy functional (an upper bound on sensory surprisal) [26]. We build on this theory to describe body illusions being driven by uncertainty minimization, i.e., the aim to obtain a maximum posterior precision or minimum posterior uncertainty. We first present the model description and notation and then detail each computational sub-process.

### 2.1. Problem formulation and parameters

Figure (1A) shows the model schematic for the body-ownership illusion based on posterior precision. It consists of the environment and the agent. The environment comprises an experimenter (E) that provides the participant (or the agent) with sensory stimuli—see Fig. (1C) for the notation used. The experimenter strokes the real hand (which is at position **Ea**) and/or the rubber hand (which is at position **Er**) locations. Thereby, the experimenter generates a force (**Et**) on the real hand, which the agent (A) can feel. Moreover, the agent can collect the visual signals **Av** and **Rv** that represent the position of the real hand and the rubber hand, respectively. The agent also feels a tactile sensation **At** on its real hand. The *y* coordinate position of the real hand **Sy** is observed by the agent as **Py** (proprioceptive measurement). Stacking these variables together we have: i) the environment state **X** = [**Ea Er Et Sy**]^*T*^ and ii) the measurements of the agent as **Y** = [**Av Rv At Py**]^*T*^.

We assume that the environment is a generative process^2^ that produces the sensory measurements from the state variables. We define this mapping as a linear function **M** between the state **X** and output (measurements) **Y** with measurement noise **z** as:

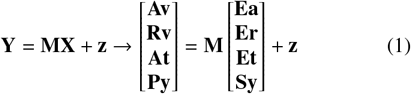

The agent processes the measurement **Y** to make sense of the world through estimation. However, the experimenter can manipulate **Y** to make the agent fall into the body illusion. We model this process within the RHI paradigm using the following components, as shown in Figure 1A: i) noise estimation (attention), ii) state estimation (body localization), iii) posterior uncertainty estimation, and iv) model selection (body ownership). The next section will explain each of these computational blocks in detail.

### 2.2. Maximum posterior precision body-ownership model

In our proposed approach, the agent selects the model that best minimises its uncertainty in estimation. This implies that the agent estimates, at each time step, both the state and the noise in the environment given the competing models or hypotheses of the perceived reality. For instance, in the RHI, the two models are “my hand is my real hand” and “my hand is the rubber hand”. In the following subsections, all subprocesses of the agent, depicted in Figure 1A, are detailed.

#### 2.2.1. Competing models of perceived reality

We assume that the brain can maintain different hypotheses of the perceived reality, where each of them defines how sensory information is generated. In the classical RHI, this accounts for two hypotheses or competing models of reality (*M*_1_ and *M*_2_). However, the this approach can be extended to any number of hypotheses. *M*_1_ accounts for accurate body estimation neglecting the rubber hand, and *M*_2_ considers that the rubber hand is our hand. We further model the participant’s sensitivity to experience the illusion by means of how both models are integrated. We define body ownership existing on a continuous spectrum parameterized by *α*. This spectrum spans between two extreme cases: a strongly non-illusory model (*α* = − 10) where the agent maintains accurate body ownership and a strongly illusory model (*α* = −10) where the agent is susceptible to incorporating external objects into their body representation. Mathematically, this is defined as the weighted combination of both non-illusory and illusory models:

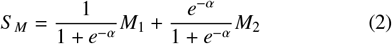

where *M*_1_ represents the mapping for accurate body perception:

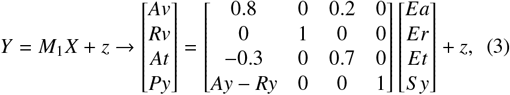

and *M*_2_ represents how sensory information will be generated if the rubber hand is our hand:

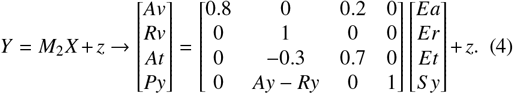

The key differences between these mappings (or generative models) lie in the processing of tactile information (*At*): while in both models tactile sensation depends on the proprioceptive estimation (*Py*), in *M*_1_, tactile sensation receives input from the real hand stimulation (*Ea*), while in *M*_2_ it depends on the rubber hand stimulation (*Er*). *M*_1_ maintains an accurate position estimation factoring in hand separation (parametrized as the distance between the hand position *Ay* and the rubber hand position *Ry*), while *M*_2_ allows information integration using the rubber hand position. Thus, both models use the hand separation parameter (*Ay - Ry*), but they apply it differently - *M*_1_ applies it to the real hand position while *M*_2_ applies it to the rubber hand position. This mathematical difference captures the essence of the illusion, where the proprioceptive signals become linked to the rubber hand rather than the real hand. Note that all variables highlighted in **bold text** belong to the generative process (environment), and the non-bold variables belong to the generative model (agent).

The inclusion of tactile force (*Et*) in our model represents the continuous nature of the stroking stimulation of the experimenter during the RHI. While *Ea* and *Er* represent the spatial locations where stroking occurs on the real and rubber hands, respectively, *Et* captures the temporal dynamics of the applied force. Having visual and tactile signals allows us to model both the spatial and temporal aspects of visuotactile integration. The tactile force parameter enables our model to capture how the strength and timing of tactile stimulation could influence the illusion. Although this implementation is a simplification of the complex spatiotemporal patterns involved in actual RHI experiments, it provides a tractable way to model continuous sensory stimulation.

##### Algorithm 1

Model selection by uncertainty minimization

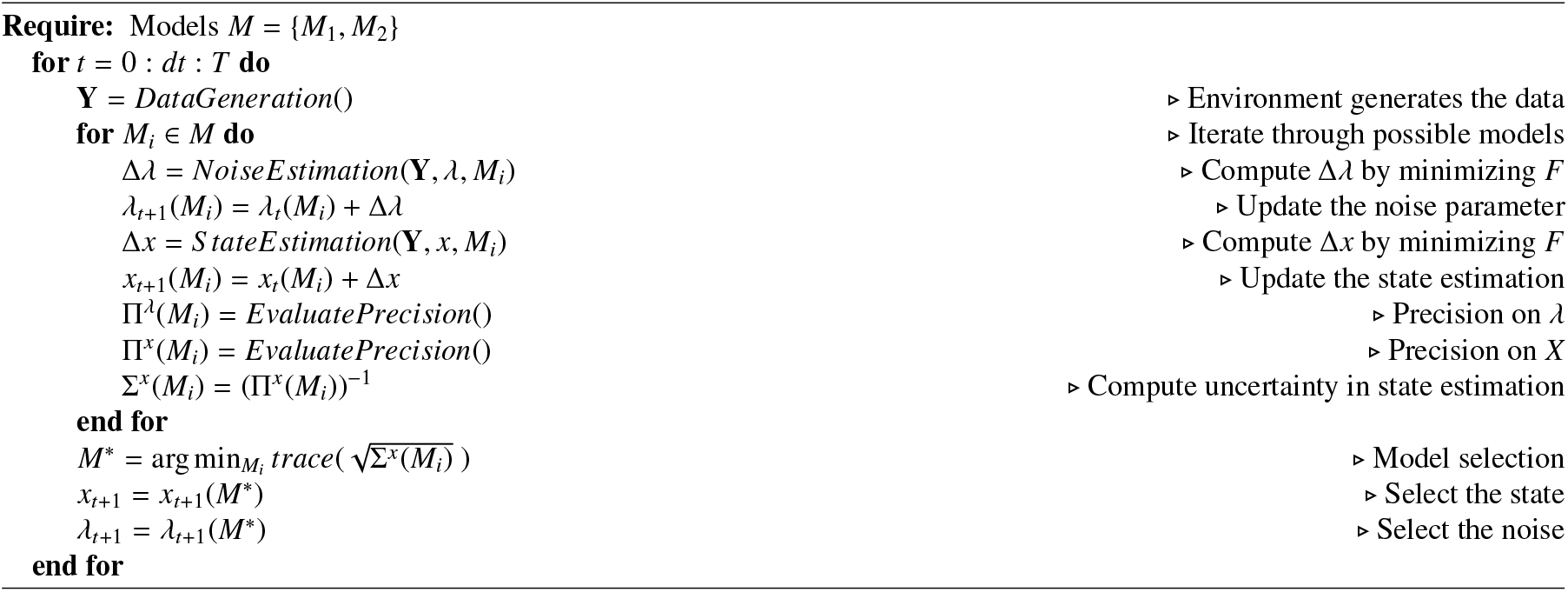

The *α* parameter of Eq. 2 determines the relative weighting between the two running hypotheses, with more negative values of *α* leading to stronger illusory effects and more positive values maintaining accurate body representation. This formulation allows us to capture both the gradual onset of the illusion and individual variations in susceptibility while maintaining the key computational features that enable both accurate body representation and illusory perception when appropriate. A participant with *alpha* = 10 will consider the information from the experimenter touching the real hand and also the force applied to the real hand as the most important, compared to the other extreme participant (*α* = − 10), which values the rubber hand as the main source of these stimuli, slightly ignoring the force *Et*.

#### 2.2.2. Model selection by uncertainty minimization

To determine the most plausible hypothesis, our agent selects the one with higher posterior precision. First, we describe the overall mechanism of model selection^3^ that produces the body ownership illusion—depicted as a pseudocode in algorithm 1—and then we detail the different computations, such as noise precision estimation, body state estimation and uncertainty estimation. The code and the parameters for replication of the results and figures can be found in Appendix 5.3.

For all competing models *M*_*i*_, we compute the posterior precision of states Π^*X*^(*M*_*i*_), along with the estimated state (*X*). We define the model selection criterion, using the posterior precision of state estimation, for the competing models as:

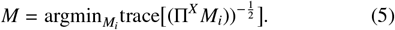

The core idea is that the agent selects the model that minimises the uncertainty in state estimation, mathematically described as the trace (sum of diagonal elements) of the square root of the posterior covariance matrix (inverse of precision matrix, Σ^*X*^ = (Π^*X*^)^−1^) of state estimation. According to the proposed model, the RHI will begin at the moment where *M*_2_ has lower uncertainty, compared to *M*_1_. Once the RHI starts, the distinction between the actual hand and the rubber hand is hypothesised to begin to fade from the agent’s point of view, resulting in a flow of information from external inputs on the rubber hand (*Er*) to the sensory perception about the actual hand (*At* and *Py*). This can be observed by comparing the differences in the last two rows of equations (3) and (4): As the agent switches from model *M*_1_ to model *M*_2_ the RHI begins.

##### Free parameters

The core free parameter of the model is the estimate of the noise parameter *λ*, which is learned dynamically through precision adaptation. This parameter captures the key phenomena. Our model has other parameters: *i*) The prior noise precision parameter *P*^*λ*^ for each sensory input; *ii*) The model matrices *M*_1_ and *M*_2_ that define how states map to observations; *iii*) The prior mean and precision (*η*^*λ*^ and *P*^*λ*^) for *λ* that helps to embed prior information for precision adaptation; and *iv*) The learning rates *k* for state and precision estimation.

#### 2.2.3. Role of precision in the computational model

In our computational model, we consider three sources of variance, i.e. three precision parameters that need to be estimated: *i)* noise precision Π^*z*^, that is parameterized using *λ* (details in appendix 5.2), *ii)* prior precision on estimates *P*^*λ*^, and *iii)* posterior precision on estimates Π^*X*^ and Π^*λ*^. While the noise precision (Π^*z*^) is learned online by our model using the agent’s prior precision (*P*^*λ*^), the posterior precision (Π^*X*^) is computed alongside the state estimates *X*. The noise precision learning contributes to the identification of the noise levels in the sensory data, while the posterior precision computation contributes to the decision (model selection) of which model to rely on (*M*_1_ or *M*_2_). The following sections will detail the mathematical formulations behind these processes.

#### 2.2.4. Joint State and Noise Precision Estimation

We model body state estimation as inferring the internal states of the body from the observations *Y*, given a model of reality. The agent keeps track of the states *X*, the activities of the experimenter *Ea, Er, Et* and the perceived hand position *S y*. This estimate is computed in the agent by approximate Bayesian inference through the minimization of the free energy [53]. The free energy of the agent that tries to predict the outputs **Y**, under state *X* and model *M*, and learns (or estimates) the measurement noise parameter *λ*, with prior mean and precision *η*^*λ*^ and *P*^*λ*^ respectively, can be written down as (see [51] for full derivation):

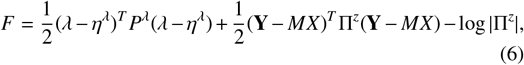

where the operation ^*T*^ denoted the transpose. Here, the noise precision Π^*z*^ is parametrized using *λ* (see appendix 5.2). The agent performs state estimation under a selected model using free energy minimization:

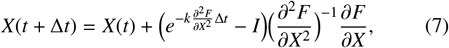

where *I* denotes the identity matrix. The update rule uses the first two gradients of *F* in Eq 7, which can be calculated by differentiating Eq (6) with respect to *X*. This mathematical routine updates *X* such that it minimises *F* in Eq 6 with respect to *X*. The agent performs online noise precision learning by updating *λ* such that it minimises the free energy, using its first two free energy gradients:

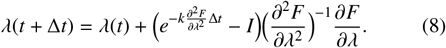

The gradients of *F* can be calculated by differentiating Eq (6) with respect to *λ*. Note that the update of *λ* changes the learned observation noise Π^*z*^ as per Eq (10) in Appendix. The updated parameter *λ* is used to compute Π^*z*^ in Eq (6). This is further used in the state estimation in Eq (7) in the next time step. Therefore, the noise parameter *λ* directly influences the state estimation. In this particular setting when the noise has a large amplitude, the agent can perceive illusory body state estimation.

#### 2.2.5. Uncertainty in estimation

According to FEP, the posterior precision of the estimate (the second-order statistic reflecting the confidence about the estimates [52]) is obtained using the second-order gradient of *F*. We use this to evaluate the uncertainty about the estimates by taking the inverse of the posterior precision of the state estimates as [51]:

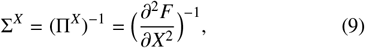

where Σ^*X*^ denotes the agent’s uncertainty in making predictions about the states, given the observed data. The agent computes the precision for both the competing models Π^*X*^(*M*_1_) and Π^*X*^(*M*_2_) to decide. It is used directly in the model selection criterion given in Eq (5). The core idea is that the agent decides to believe in a model that best minimizes (or resolves) its uncertainty in estimation. The hypothesis is that once the visual signals of the hand (*Av*) are highly noisy (hand covered), the illusory condition 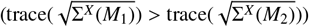 becomes true and the agent switches its model selection from a non-illusory model (*M*_1_) to an illusory model (*M*_2_), resulting in the experience of RHI.

### 2.3. Evaluative measures for the model

Now that it has been explained how the agent uses the uncertainty in state estimation for model selection, we conclude with a description of how to model hand coverage, the illusion sensitivity and measure proprioceptive drift. Note that these measures are not modeled as part of the agent in Figure 1, but are used as fitted parameters for evaluations and comparisons.

#### 2.3.1. Modeling hand coverage

We model the real hand coverage by making the visual signal very noisy. The measurement noise **z** is composed of four components (**z**^**Av**^, **z**^**Rv**^, **z**^**At**^, **z**^**Py**^). When the real hand is covered, the visual signal **Av** obstruction is modeled by adding a high noise **z**^**Av**^. Analogously, for no coverage of the real hand, we use a low noise **z**^**Av**^ on the signal. In broad terms, the noise level **z**^**Av**^ modulates the error in the visual information input. This modeling, although controversial, is grounded on the idea that darkness is completely uncertain [30] and hence, that the absence of visual information invokes a strong noise component—see discussion (Sec. 4.3) for further explanation.

#### 2.3.2. Individual differences through illusion sensitivit

There is a high variability between individual susceptibility to the body illusion. Moreover, depending on the manipulation (e.g., temporal synchrony), the body illusion sensitivity varies [45]. To account for the agent’s sensitivity to experience the illusion, we modulate the *α* parameter from Eq 2. Technically, reliance on prior versus posterior evidence of each participant is modeled through parameter *α*. Intuitively, the higher the *α*, the less susceptible the agent is to experience the illusion. Thus, Eq (2) captures the variability of body ownership sensitivity, which also influences the experienced proprioceptive drift.

#### 2.3.3. Measuring the proprioceptive drift

During the RHI experiment, the agent’s state estimate of it’s hand position *S y* may be disrupted, due to the high noise amplitude in the visual data (**z**^**Av**^), leading to the experience of a proprioceptive drift, where the perceived hand position *S y* shifts from the real hand position *Ay*, towards the rubber hand position *Ry*. The agent cannot report the proprioceptive drift during the RHI because it is an illusion, and the agent does not have access to the true signal **Sy**. The proprioceptive drift in this paper is computed by taking the difference between the real position of the hand **Sy** and the agent’s estimated hand position *S y* as shown in Figure 1.

## 3. Results

Here, we show using simulations how the proposed precision-based model may explain the mechanism and dynamics behind body ownership illusions and their relation to multisensory integration and localization drifts. To this end, we simulated the RHI paradigm, being one of the most paradigmatic body perceptual manipulations, where a participant experiences the illusion that an isolated fake hand becomes an agent’s hand. The parameters used to generate the results are described in Appendix 5.3. We describe the following results:

1. We validate the proposed model in a simulated RHI experiment (Fig. 2), where we show how posterior uncertainty minimization under the two competing models of reality yields to a perceptual switch that produces the illusion of owning the fake hand.
2. We analyse the precision adaptation mechanism of the proposed model, which is underneath the perceptual switch (Fig. 3). The introduction of visual noise by covering the hand is one of the factors producing the effect. The online adaptation to the sensory noise through precision estimation appears as an essential property to experience the illusion.
3. To further validate our model, we show that breaking the noise precision learning – where the model does not adapt to the sensory noise – leads to a state where switching the current model of reality to a different one is implausible, hence, no illusion can occur (Fig. 4).
4. We assess the dependence of the body ownership illusion (model switch) on the distance between the rubber hand and the participant’s own hand (Fig. 5)
5. We evaluate how the proprioceptive drift depends on the selected competing model (Fig. 6), where *M*_1_ produces noisy estimations but not drift, while when selecting *M*_2_, the model produces the experimentally observed proprioceptive drift.
6. We validate the effect of the inter-hand distance on the proprioceptive drift dynamics, given that the illusion is happening (Fig. 7).
7. Finally, using the modeling of artificial participants with individual differences, we explore the effect of the participant’s susceptibility to experience the RHI (*α*) and the inter-hand distance (*Ry*) on the generated proprioceptive drift (Fig. 8). The results show a similarity between the profiles found in a recent experiment in humans and macaques [22] and our model.

**Fig 2:**
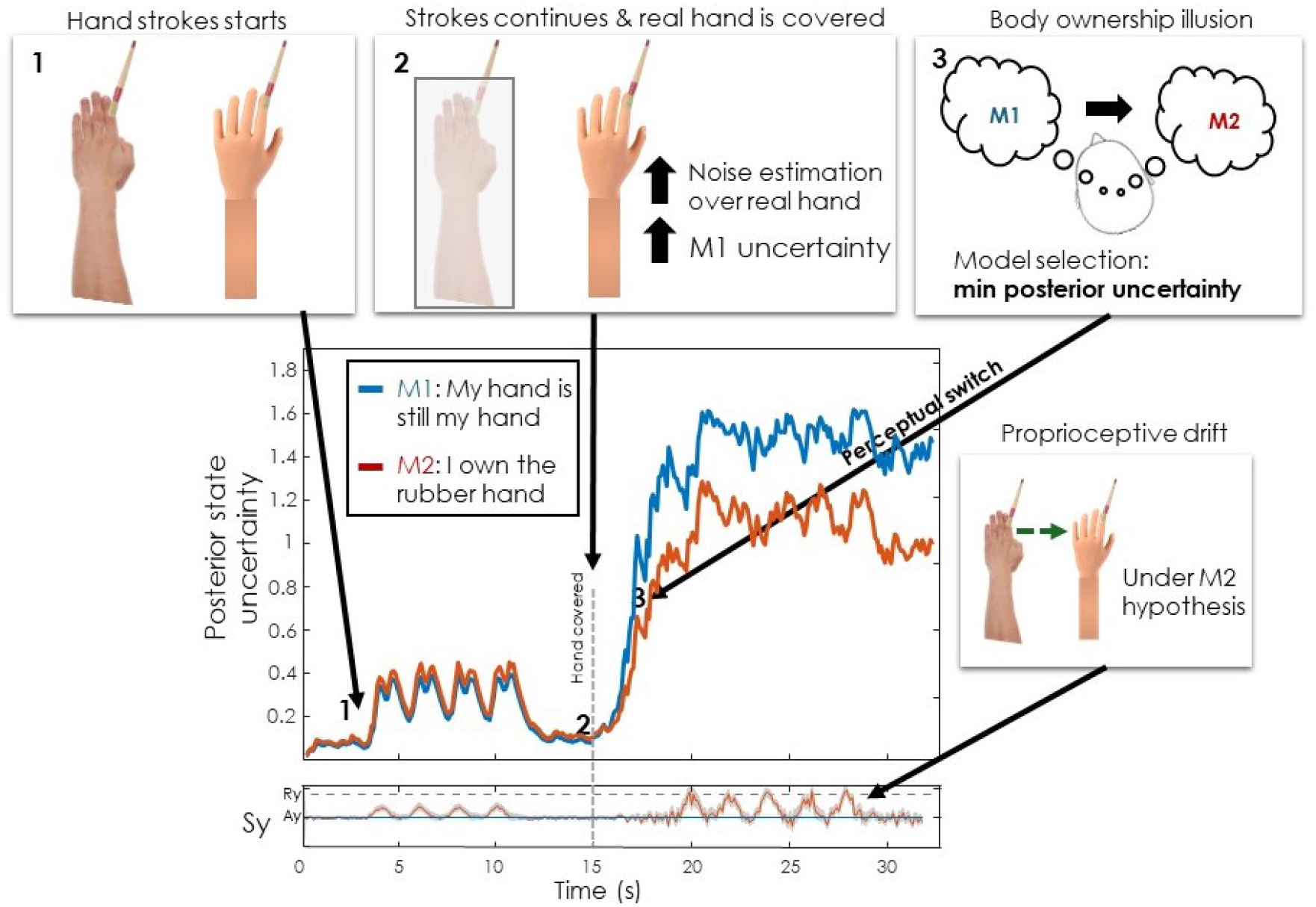
Body-ownership perceptual switch. The experiment sequence and its results in our model: 1) Both hands are being simultaneously stroked; 2) The real hand is covered and the strokes continue; 3) The participant starts experiencing the RHI. In the bottom graph, the competition of the two models and their uncertainty during the experiment are shown. The middle panel shows the estimated posterior state uncertainty ((Π^*X*^)^−1^) of both models and the bottom panel the perceived position of the hand (*S y*) given the participant believes in model *M*_1_ (blue) or in model *M*_2_ (red). After the hand is covered, there is a switch between these two models, as *M*_2_ starts being more precise (lower uncertainty). This then leads to believing that the rubber hand is the participant’s hand, thus leading to a proprioceptive drift depicted in the bottom graph. Note that the blue line in this panel represents the real proprioceptive signal *S y*, whereas the red line represents the estimated *S y* under the assumption of *M*_2_. In this simulation, we used rubber hand position *Ry* = 0.8.

**Fig 3:**
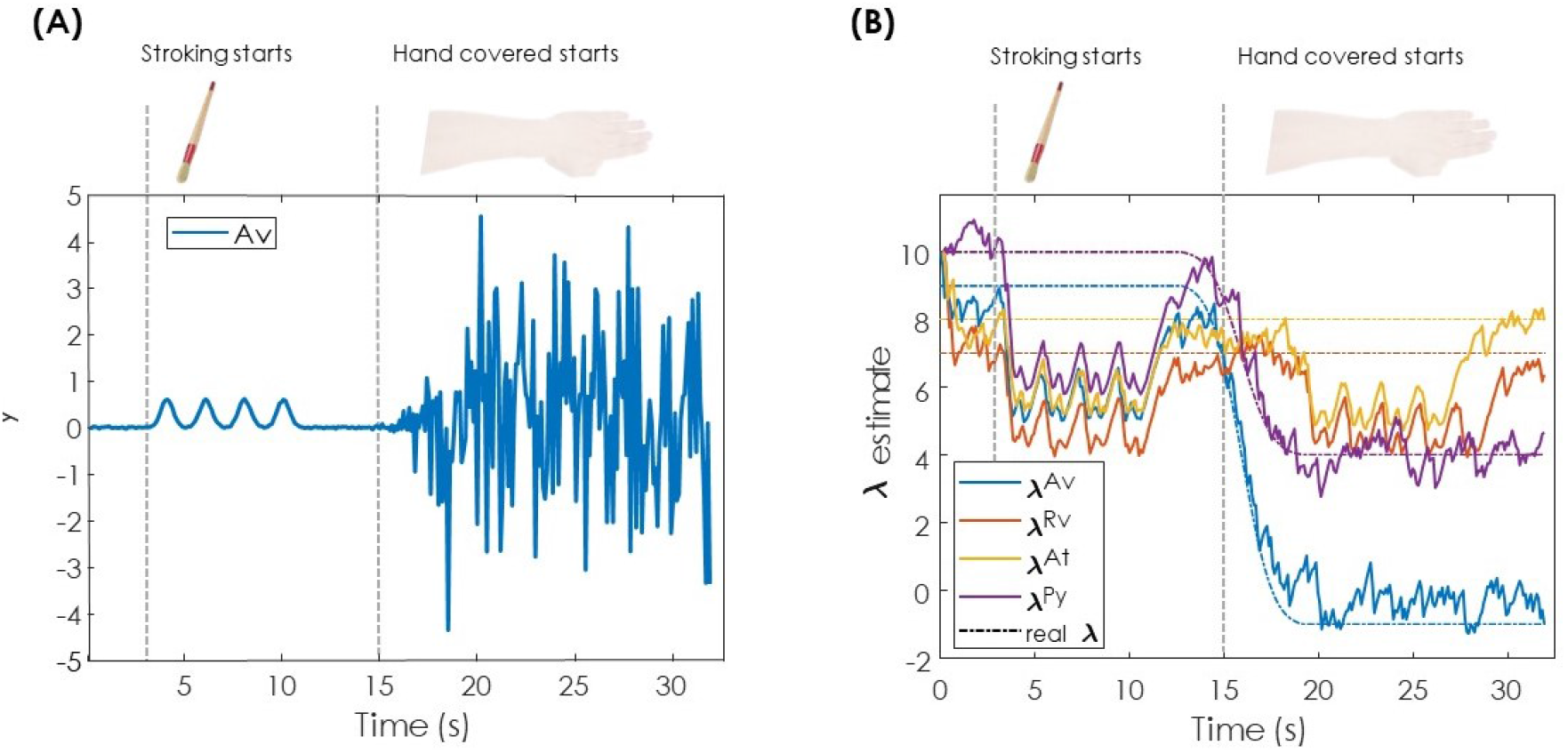
Precision adaptation (attention). **(A)** Visualization of visual signal from the real hand *Av* signal in blue. It is highly noisy after the real hand is covered (at time = 15 seconds). This represents the highly uncertain nature of the visual data for the hand position once it is covered.**(B)** Noises estimate block from our model in Fig. 1, given the sensory measurements. The colored plots track the true noise parameter *λ* (in dashed coloured lines), showing that our model is capable of correctly estimating the precision levels of all sensory signals. In other words, the uncertainty of the sensory signals is fully captured by our model in real time. Simulation parameters can be found in Appendix 5.3.

**Fig 4:**
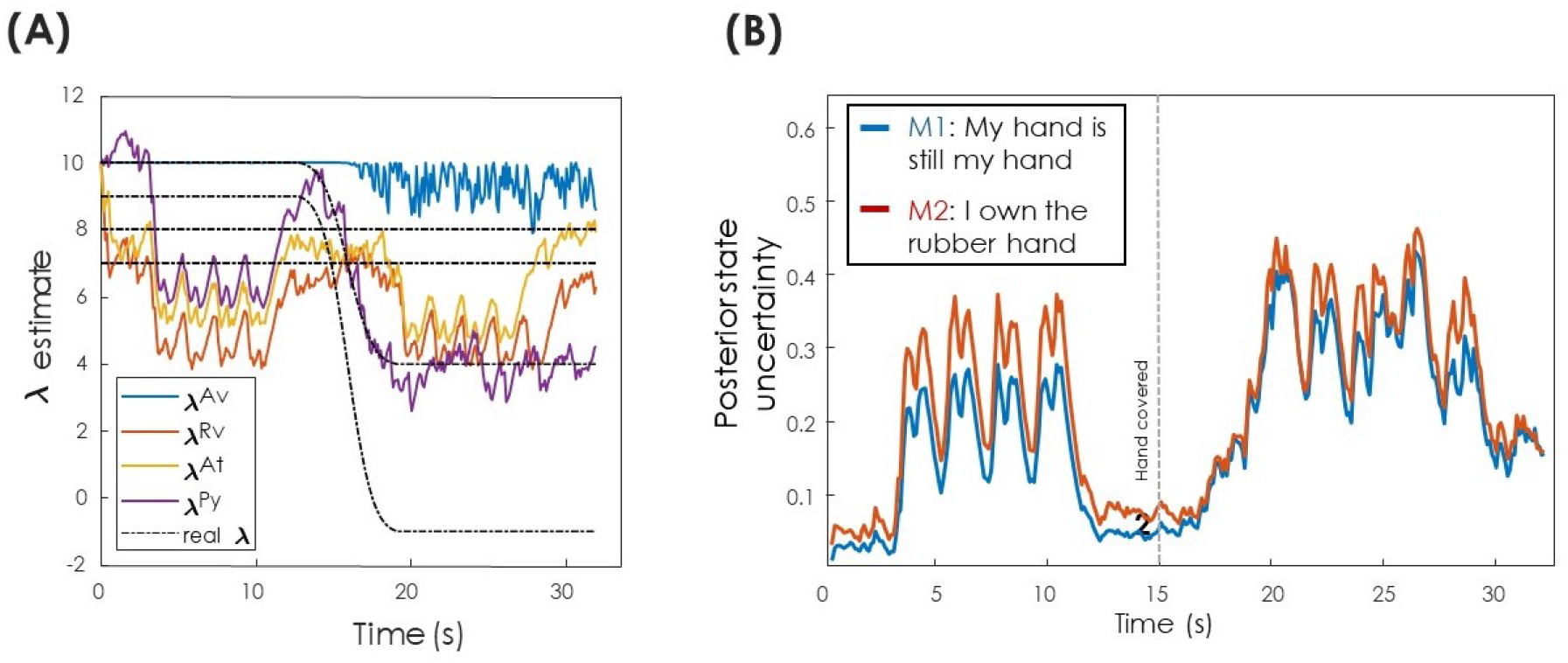
Breaking precision adaptation leads to no illusion. **(A)** This figure is similar to Figure 3B, except for the fact that the agent is now biased towards believing that the signal **Av** is highly precise by using a higher prior precision (*P*^*λ*^ = diag([*e*^10^, *e*^−4^, *e*^−4^, *e*^−4^])) for the signal *Av*), although it is as noisy as in Figure 3A. Note that *λ*^*Av*^ (in blue) stays high, around 10 instead of tracking the real *λ* (in black). **(B)** If an agent is (incorrectly) overconfident over the visual signal Av, it will lead to a behaviour where the two competing models will not switch—the red curve always stays above the blue one (compare with the switching in Figure 2). Simulation parameters can be found in Appendix 5.3.

**Fig 5:**
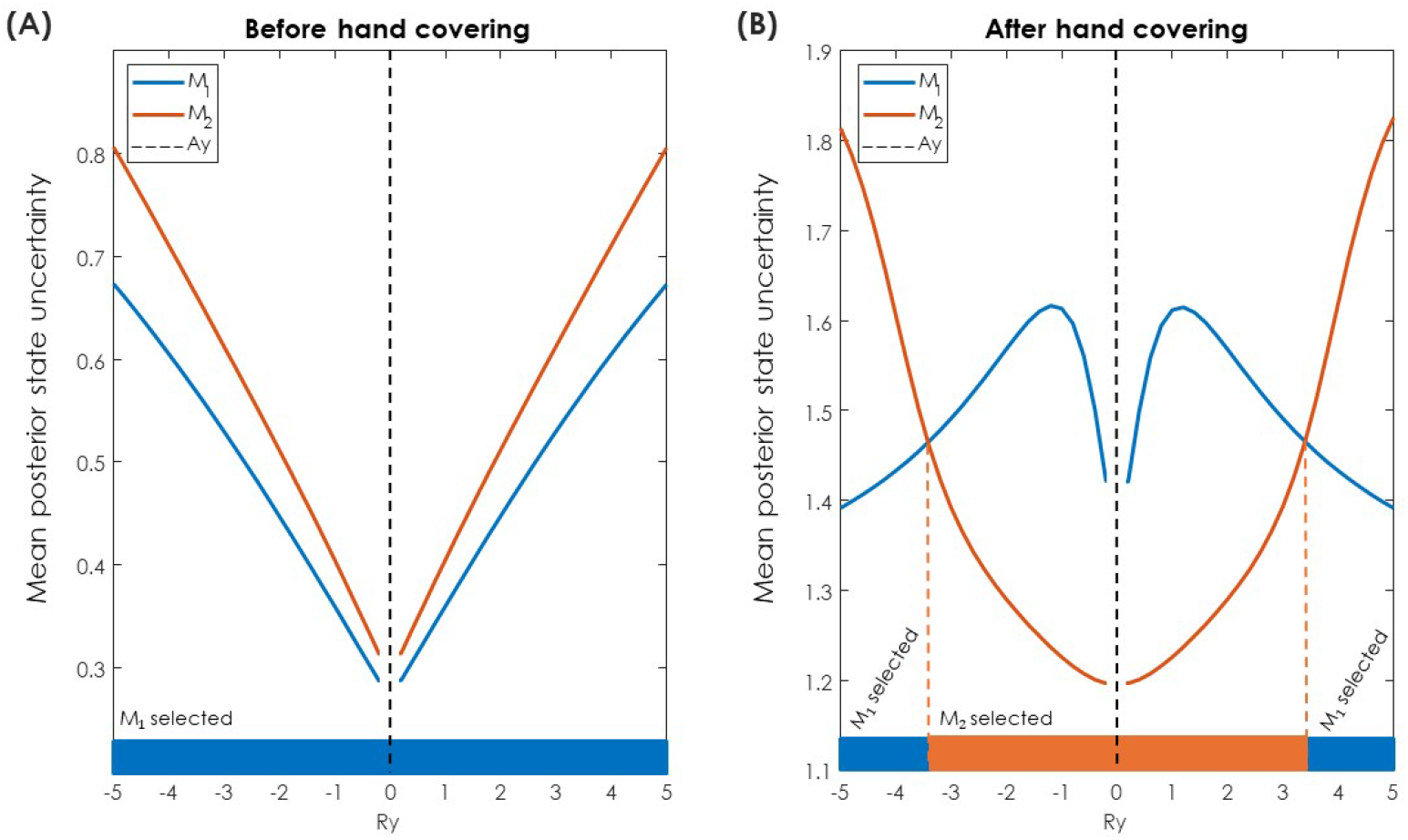
Model switch is dependent on the inter-hand distance. The vertical axis of panel A represents the mean posterior state uncertainty of all 4 signals (**Ea, Er, Et, Sy**) averaged when hand is uncovered (from the 4th to the 11th second), while in panel B, the average when hand is covered is from 20th second to the 30th second. These averages are then plotted for both *M*_1_ and *M*_2_ for different inter-hand distances. The horizontal axis represents different positions of the rubber hand (*Ry* = − 5 : 0.2 : 5). Notice that the value of *Ry* = *Ay* = 0 was removed, as it would mean that the fake hand is literally lying on the participant’s real hand. The colors on the bottom indicate whether the model remains at *M*_1_ or switches to *M*_2_ under a given *Ry*. Simulation parameters can be found in Appendix 5.3.

**Fig 6:**
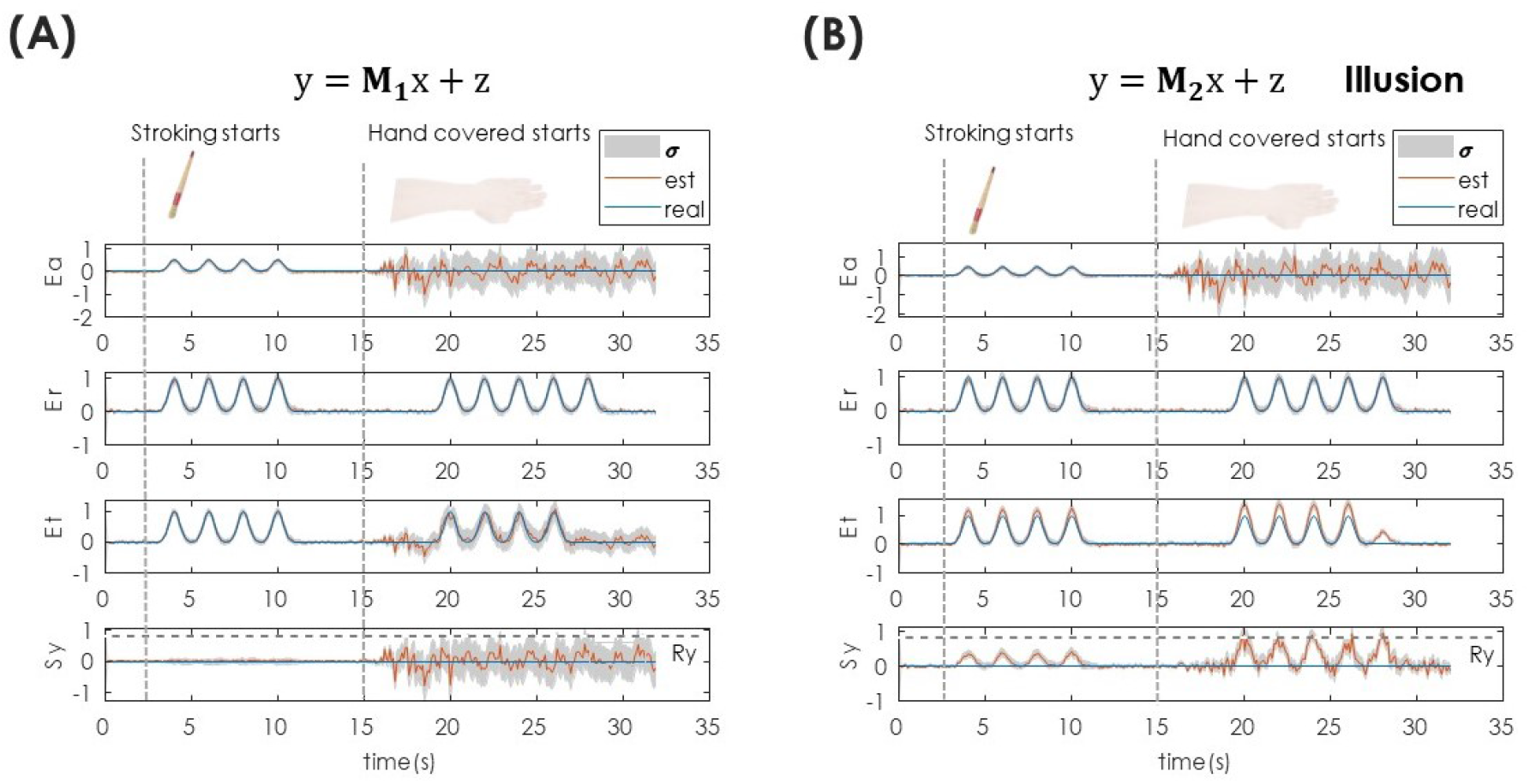
Replicating the proprioceptive drift. **(A)** State estimates of all the variables under the non-illusory model *M*_1_. **(B)** The same state estimates when the illusory model *M*_2_ has been chosen. Both figures show state estimates of all input data *Ea, Er, Et*, and *S y*. Only in the illusory model on the right, the proprioceptive drift (*S y*) and the related touch (*Et*) are observed. Also note the increase of noise in *Ea* in under both models when the hand is covered. The distance between the rubber hand and real hand is 0.8 in this case. The distance dependence on the proprioceptive drift is evaluated in Figure 8.

**Fig 7:**
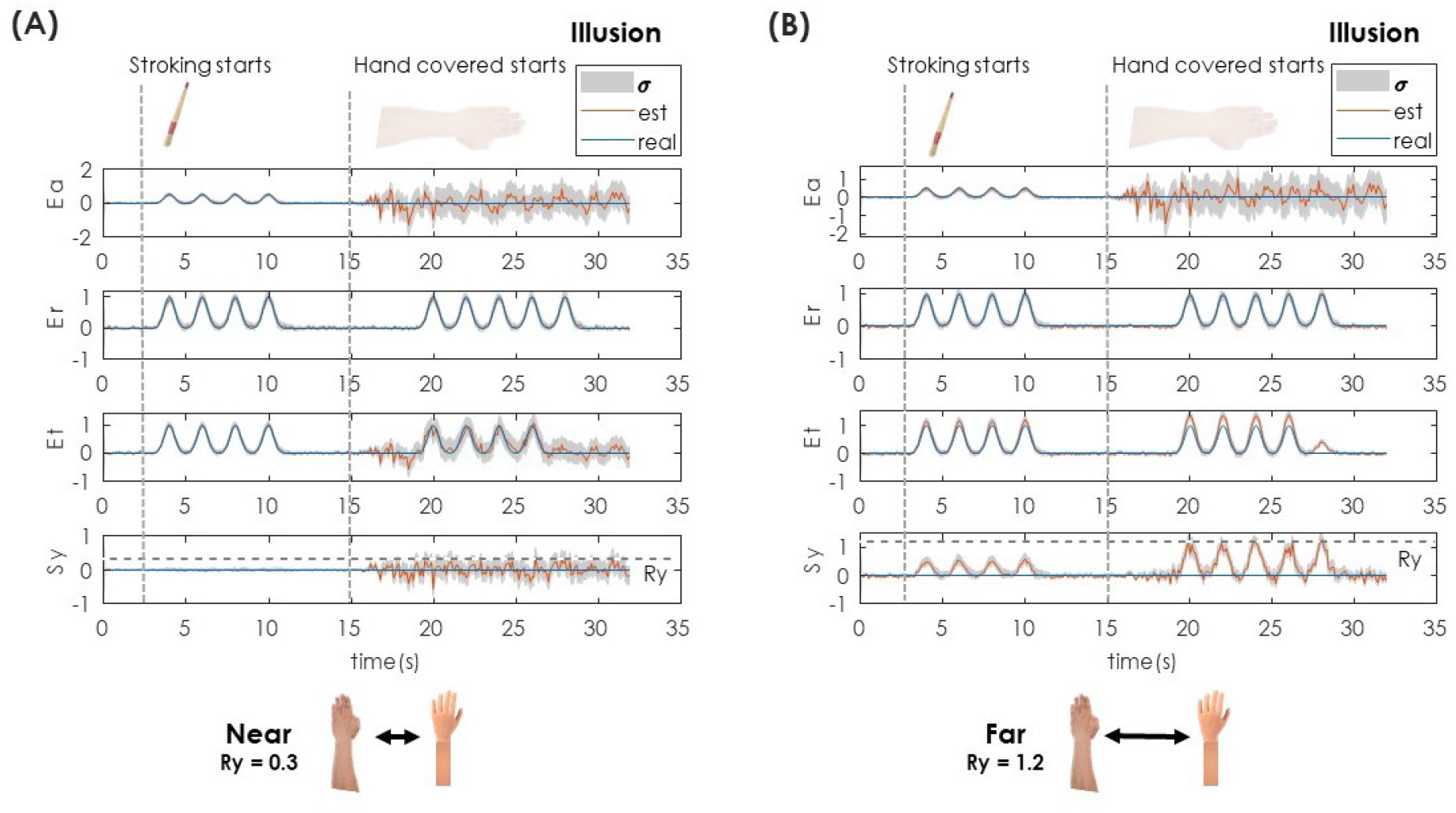
Inter-hand distance influence on the preceptual drift. The same template as in Figure 6 was used. Here, however, both figures represent the state estimates under the illusory model, where we varied the interhand distance. **(A)** Near (*Ry* = 0.3). Effect of *M*_2_ when the inter-hand distance is small (Ry = 0.3). **(B)** Far (*Ry* = 1.2). Effect of *M*_2_ when inter-hand distance is increased. This shows that with the increase of the distance between the hands, the proprioceptive drift becomes stronger (or increases in amplitude).

**Fig 8:**
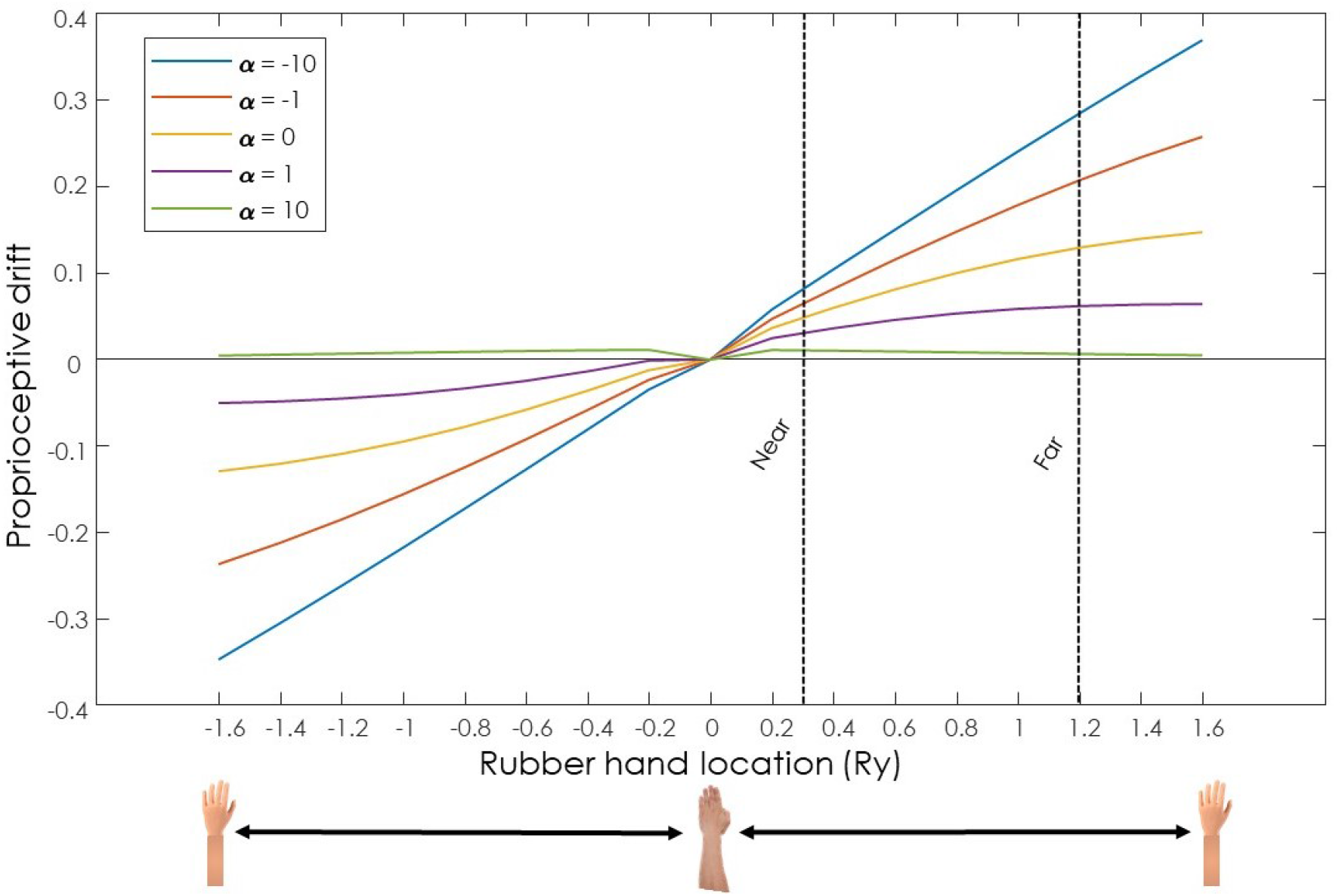
Dependence of the inter-hand distance and the participant’s susceptibility to experience the illusions on the proprioceptive drift. Inspired by [22], we show a similar trend from previous empirical findings where the modulation of the inter-hand distance and different subjective experience to the illusion affects the proprioceptive drift. Every line in the plot can be related to individual differences when transiting from *M*_1_ non-illusory to *M*_2_ illusory models (i.e., susceptibility to experience the illusion). This is modulated through the parameter *α*. Here, *α* = 10 and *α* = − 10 are equal to models from equations (3) and (4), respectively. We measured the proprioceptive drift as the average of *S y* between the 17th and the 30th second. Note that the dashed lines represent the distances for near and far conditions from Figure (7).

### 3.1. Body-ownership illusion: model selection based on the uncertainty minimization

Figure 2 describes the proposed model behaviour, as described in algorithm (1), as a function of uncertainty estimation when the variance of the sensory input increases (Fig. 2). First, the posterior uncertainty ((Π^*X*^)^−1^)—when both hands are visible and being stroked by the researcher (Figure (2, panel 1)—favors the non-illusory hypothesis (*M*_1_). Once the hand becomes covered (Figure 2 panel 2), there is an immediate increase in *M*_1_ uncertainty, which is due to the increase of the noise estimate of the real hand (*λ*). After a few moments, there is a switch between the two models regarding posterior uncertainty (Figure 2 panel 3). This leads to the participant believing that they own the rubber hand, as the *M*_2_ hypothesis now provides a lower uncertainty compared to *M*_1_. As this is happening, we can see that in model *M*_2_, there is also an observable proprioceptive drift driving away from *Py*=0 towards the location of *Ry*=0.8 (red line, bottom panel). Therefore, this simulation shows the two main effects of body-ownership illusions and the potential computational mechanisms responsible for body ownership (minimizing uncertainty) and proprioceptive drift or mislocalization (state estimation as multisensory integration).

### 3.2. Estimating noise through precision adaptation

We evaluated the reliability of our model when estimating the sensory noise. Figure (3) shows that both proprioceptive (purple line) and visual (blue line) signals from the real hand become noisy after the hand is covered at the 15th second. This is because covering the real hand results in noisier visual and proprioceptive signals. The coloured lines in Figure 3B follow the real noise levels (the dashed lines) in all signals, showing that model is capable of correctly estimating the precision levels of all sensory signals from data.

### 3.3. Validating the model via breaking precision adaptation

We hypothesised that precision learning/adaptation is a necessary precondition for model switching. In other words precision learning is crucial for the model switching, and if this is broken (by providing priors that inhibit learning), the switch will not occur. To test this, we disrupted the noise precision learning by biasing it with a high prior precision on noise. We defined (*P*^*λ*^ = diag([*e*^10^, *e*^−4^, *e*^−4^, *e*^−4^])) for the signal *Av*. In Figure (4), the effects of this manipulation are shown. Figure (4A) shows the improperly learned precision signals of the input *Av* due to the high prior on *λ*. Consequently, Figure (4B) demonstrates that the posterior state uncertainty (Π^*X*^)^−1^ of *M*_1_ remains consistently lower than *M*_2_, regardless of hand coverage in our earlier findings in Figure (2). These results provide face-validation for the necessity of precision learning/adaptation in achieving the body-ownership illusion under our proposed model.

### 3.4. Dependence of the illusion on inter-hand distance

We further examined how the illusory perceptual switch depends on the distance between the real and the rubber hand (|*S y* −*Ry*|). Empirically, the RHI breaks down when the fake hand is too far from the real one, as the brain struggles to integrate highly discrepant multisensory information. Our model captures this effect naturally (Figure 5). Because *M*_1_ has a lower uncertainty with an increased inter-hand distance, the illusory state cannot emerge. Figure 5A shows that the non-illusory model *M*_1_ is always more certain about the states for the time frames when the is not covered, independent of the distance *Ry*. Figure 5B shows the model’s posterior state uncertainty for the time frame when the hand is covered. When the rubber hand is too far from real hand (when |*Ry*| > 3), the non-illusory model becomes more certain (blue curve below the red curve), implying that the RHI will not happen when the inter-hand distance is too high. Notice that Figure 5A and B are completely symmetrical, highlighting that the model behaves in the same manner with negative and positive values.

### 3.5. Proprioceptive drift in competing models

We have shown that our model can explain the RHI as a result of adaptation to the precision of sensory stimuli. In this section, we will now show how our model explains proprioceptive drift in more detail than in Figure 2, where it is mentioned only briefly as an independent mechanism that happens alongside model selection. Here, we show that proprioceptive drift occurs only in the illusory model *M*_2_ (Figure 6B) but not in the non-illusory model *M*_1_ (Figure 6A). These plots represent a single trial of the experiment described in Figure 2. In Figure 6, the red lines that represent the estimated perception of the signal coincide with the blue lines that represent the real values of the experimenter’s stimulation. Therefore, the agent correctly estimates the location of the touch of the real hand (*Ea*) (Figure 6 first panel), of the fake hand (*Er*) (Figure 6 second panel), and the tactile sensation (*Et*) (Figure 6 third panel). However, as shown in Figure 6B, even during the first half of the experiment, when both hands are visible, the *M*_2_ model produces a small deviation in the estimated hand position *Sy* from its true value (Figure 6B in the bottom panel). Here, the red curve deviates from the blue line towards the dashed horizontal line (*Ry*). This plot highlights that the model is able to generate the proprioceptive drift.

The proprioceptive drift also occurs before the hand is covered, though participants would not experience this drift since the non-illusory model *M*_1_ has a lower posterior state uncertainty compared to *M*_2_ (see Figure 6A last panel for comparison, and Figure 2 for model selection). This demonstrates that the proprioceptive drift exists within the mathematical structure of the *M*_2_ model even before it becomes the selected model. Moreover, we show how the process of model selection is independent from the mechanism of drift computation. Only when *M*_2_ becomes the selected model after hand occlusion does this drift become part of the agent’s experienced hand position.

#### 3.5.1. Dependence of proprioceptive drift on inter-hand distance

The dependence of the proprioceptive drift on the inter-hand distance has been observed empirically in many experiments [62, 42]. Here, we also validate our model with this logic. In Figure (7), we compare two illusory models where the distance of the two (real and rubber) hands is increased. In Figure (7A), the inter-hand distance is smaller (*Ry* = 0.3) compared to the one in Figure (7B, (Ry=1.2). In the latter figure, the estimate in *Er* is incorrect, which leads to a smaller and shorter proprioceptive drift (measured as the difference between the real and estimated values of *Ry*, bottom panels) compared to Figure (7A).

Next to the inter-hand distance, we also we modulated the strength of both the illusory and non-illusory models via the parameter *α* in the softmax function in Eq (2) – see methods for more detail. By doing so, we assume that we modulate the subjective susceptibility of the competing models. We vary these two parameters, the inter-hand distance *Ry* and the susceptibility *α*, independently, to show that the proprioceptive drift is increasing in amplitude when slowly increasing the distance between the two hands. The speed of the amplitude increase becomes smaller with higher distance (Figure (8)). In this figure, a higher *α* value is associated with the agent being less susceptible to the illusion and vice versa. We plot the maximum value of the proprioceptive drift for simplification, which is represented on the vertical axis, while the distance between the subject’s and the rubber hand is on the horizontal axis. In addition, we show that modulating the subjective strength of the illusory model results in different strengths of the proprioceptive drift, as shown in [22].

In the introduction, we touched upon the discussion of whether proprioreceptive drift is a good indicator of the RHI. The results, from our perspective, also link the difference between the selection of the illusory model and feeling the drift, and not selecting the illusory model and still feeling the drift, ultimately supporting the case of two distinct computations for proprioceptive drift and illusion per se. Here, we therefore argue that these two are the result of separate but related computational mechanisms [37, 38, 67, 68] – state estimation and precision maximisation.

## 4. Discussion

We have proposed a novel theoretical model that can explain body illusion dynamics and evaluated it through a simulated RHI experiment. We built a general model that selects between competing interpretations of body representations and switches between them by maximizing the precision (minimizing the uncertainty) of the state estimation (Fig. 2). The model was face-validated on previous experimental results via replicating the proprioceptive drift (Fig. 6), its dependence on the inter-hand distance (Fig. 7), and the illusion dependency on the inter-hand distance (Fig. 5). We showed that the model can reproduce the observed differences in proprioceptive drift among participants due to the subjective susceptibility towards illusion—modulated by the *α* parameter (Figure 8). Overall, our model provides *i*) useful insights into the process dynamics during the RHI, going beyond previous static computational models, and *ii*) a unified mathematical account that can distinguish between the ownership illusion and the perceptual drift.

In the following subsections, we discuss how this model’s predictions may contribute to future research in body ownership: its neural underpinnings, its relation with other recent body-ownership models, and how to empirically test the model with the challenges and limitations of the current implementation.

### 4.1. Potential neural underpinnings of the model

The current work introduces the model and does not explicitly embed the computations in a specific brain region. However, some neurobiological clues exist regarding where such computations could be performed in the brain. Recent neuroimaging evidence points to a network of regions involved in body ownership illusions, particularly the premotor cortex and posterior parietal cortex (IPS) [10, 31]. The posterior parietal cortex is especially relevant to our precision-based model, as it plays a key role in multisensory integration and uncertainty processing. Recently, Chancel et al. (2020) [8] directly applied a causal inference model to fMRI data, identifying the posterior parietal cortex as crucial for resolving uncertainty during body ownership decisions. Additionally, the premotor cortex shows consistent activation during body ownership illusions [17], suggesting its involvement in maintaining coherent body representations under sensory uncertainty.

### 4.2. Alignment with other uncertainty-based models

There is a strong connection between our model and other models built on Bayesian Causal Inference [71]. However, there are important differences. For instance, Chancel and colleagues [10] defined a model where the implementation of uncertainty relies on visual noise manipulation via augmented reality with three discrete noise levels. However, our model treats uncertainty as a continuous variable that dynamically adapts through precision estimation. This fundamental difference in uncertainty handling reflects distinct theoretical approaches to the decision-making process. In [10], a fixed decision criterion is used for each noise level to study how participants infer common cause probability, while our model employs online precision adaptation, where the system actively learns about and adjusts to sensory uncertainty. These models should be viewed as complementary rather than competing accounts. Relevantly, [10] provides rigorous psychophysical evidence for the role of Bayesian inference in body ownership, demonstrating how increased visual noise facilitates ownership illusions. This aligns with our model’s predictions about the relationship between sensory uncertainty and integration. However, our model extends this by suggesting how the brain might continuously update its estimates of sensory precision rather than simply responding to externally manipulated uncertainty levels. Their careful experimental paradigm validates key principles that our more mechanistic model assumes, particularly regarding how the brain weighs and integrates multisensory information under varying levels of uncertainty.

The combination of Chancel et al’s experimental rigor [8, 10] with our more mechanistic model of precision estimation suggests promising directions for future research. Their findings about the relationship between synchrony detection and ownership judgments could be extended to test conditions of continuously varying uncertainty, as predicted by our model. Similarly, our model’s predictions about dynamic precision estimation could be tested using modified versions of their psychophysical paradigms. Together, these approaches could provide a more complete understanding of how the brain determines and maintains body ownership through precision-weighted integration of multisensory signals.

### 4.3. Testing our model and limitations

Our computational model of body ownership based on precision adaptation makes several testable predictions that could be empirically validated. Here, we outline experimental approaches that could confirm or challenge key aspects of our model, including *i*) predictions about dynamic proprioceptive drift, *ii*) the mechanisms of model selection versus averaging, the distinction between uncertain and absent signals, and the temporal dynamics of the illusion. These proposed experimental tests would help validate our theoretical framework and potentially refine our understanding of the computational mechanisms underlying body ownership illusions.

#### Dynamic proprioceptive drift prediction

A key prediction from our precision-based model concerns the temporal dynamics of body ownership illusions. Unlike static models that focus only on end states, our approach captures moment-to-moment changes in the proprioceptive drift (PD). Our model thus predicts that the PD should dynamically vary with each brush stroke during the RHI induction rather than being a static effect measured only after the illusion. While current experimental paradigms typically measure PD only before and after the induction period, this prediction suggests the need for continuous measurement techniques to capture potential stroke-by-stroke variations in perceived hand position. Such dynamic measurements could provide crucial validation of our model and offer new insights into the temporal dynamics of body ownership illusions. There are already several methodologies that can be directly implemented in the model [66, 43], awaiting further tests of the precision-based theory developed here.

#### Model selection versus model averaging

Although we used model selection as the mechanism behind for reporting body ownership, we also offered a way to perform model averaging through the weighted combination to capture individual differences and the illusion sensitivity. While model averaging is the most widely accepted computational account to combine different uncertain models in the brain, we took inspiration from the affordance competition hypothesis (ACH) that states that there are at least two competing neural correlates for distinct actions in the parieto-dorsal stream [12, 11, 70]. Once a decision is made, only one such neural activation of hand position can remain. This leads to the final action, referred to as the winner-take-all algorithm. The ACH slightly resembles our modeling effort, but there are differences. Our approach does not consider a ‘simple’ hand movement but rather a more complicated model that combines body ownership via an intricate network of multisensory integration. Thus, different models of body ownership compete rather than actions that are generated from these models. Ultimately, though, we can consider a similar switch in neuronal activity of brain regions as characterized by the ACH showing similar neuronal patterns.

#### Uncertain signals versus no signals

One of the assumptions of our body-ownership illusion model states that although the real hand is “invisible”, it still produces a signal that is highly noisy – from the agent’s perspective – rather than not producing a signal at all. This plays a role in the switch between the two models and can be used to study different levels of perceptual manipulation where, instead of fully hiding the hand, the degree of visibility is manipulated (similarly to [10]). We emphasized that the primary driving mechanism stems from the adaptation to the changing precision of the sensory input signal that triggers the switch between hypotheses or models of reality. Assuming high noise of an invisible hand rather than no signal is an alternative way of modeling input signals, which is, however backed up by logical arguments regarding the free energy principle called the dark room problem [30]. Simply, if the goal of an agent is to resolve all uncertainty, how come all agents do not live in a dark room, where everything is completely expectable? The resolution is that darkness is completely uncertain.

#### Time scales, proprioceptive drift and onset illusion

In our simulations (Figures 6 and 8) the proprioceptive drift appears to follow the touch of the stimulation and dissipates when the strokes no longer occur. This, in empirical settings, would result in a participant perceiving the proprioceptive drift only during an episode of touch. A more stable belief of the proprioceptive drift can be obtained using temporal integration, operating over multiple timescales, such as implementing a slow-updating belief state that retains information over 10-30 seconds rather than the rapid updates (sub-second) currently implemented in our model, for instance, involving exponentially weighted averaging of the proprioceptive position estimate across multiple touch episodes, similar to integrating over the signal across a longer period (as in [35]).

Another issue related to time scales within the simulation is that our model predicts an unrealistically rapid onset of the ownership illusion and proprioceptive drift. While empirical studies show the rubber hand illusion typically requires 10-30 seconds to develop [8, 19, 41], our model shows effects within 2-3 seconds of hand occlusion. This discrepancy likely stems from our simplified implementation of precision adaptation dynamics. In reality, the brain may require longer periods of sustained multisensory evidence to adjust precision weights and switch between competing models of body ownership. Incorporating slower learning rates or additional constraints on precision adaptation will better match the empirically observed temporal dynamics. Additionally, rather than implementing a binary switch between hypotheses, a more gradual accumulation of evidence through probabilistic model averaging might better capture the progressive emergence of the illusion reported in experimental studies.

#### Temporal synchronicity

While we model synchronicity generation of the signals within the participant’s sensitivity [45] to the body illusion through the *α* parameter, one of the main limitations of our model is that the simulation does not account for deviations in the visuo-tactile temporal synchronization of the strokes, a well-studied aspect of the RHI. Thus, explicit temporal synchronisation of the signals should be incorporated to allow fair comparison with human data experiments.

### 4.4. Future work

In summary, future research should focus on the experimental validation of the model to assess the dynamics of switching between competing models depending on the posterior uncertainty (precision), as proposed in our study. In this context, researchers could create scenarios where multiple states of body ownership are simultaneously induced, allowing only one to predominate. The introduction of different temporal scales and explicit temporal synchronicity could refine the model to align with recent human experimental data, allowing for more direct empirical testing. Furthermore, the generalisation of the proposed model for any competing hypotheses would enable the study of probabilistic transition dynamics between different competing perceptions of reality. These modifications could potentially explain phenomena such as double-limb and supernumerary limb illusions [32, 21], providing valuable insights into the mechanisms of body ownership.

## Acknowledgments

FN and FZ have received funding from, Serotonin & Beyond project, the European Union’s Horizon 2020 Marie Sklodowska-Curie Actions under grant agreement No. 953327. AM is funded by the EU Metatool project (Grant agreement 101070940) under the EIC Pathfinder program. The research has been partially funded by the DEEPSELF project (467045002 DFG SPP The Active Self).

## 5. Appendix

### 5.1. Related computational models of body illusions

This section discusses previous computational models of body illusions and how they approached the topic as a problem of multisensory conflicts, especially the RHI [6, 35, 48] and its virtual reality (VR) version [73, 46]. One of the first seminal works on modeling RHI comes from [71], where a model driven by Bayesian inference is proposed, focusing on the synchronization of stimuli in time. The authors went beyond the optimal integration model [22], by introducing causal inference, where two potential models of reality are presented: either the tactile information is coming from the subject own hand or from the fake hand. However, this account presents body ownership as a static probability computation, where no explanation is given on the inner functional mechanism producing the dynamics of the perceptual drifts nor the illusion [25]. Bayesian causal inference was revisited in [10], focusing on the impact of the input noise to the emergence of the RHI.

Another line of research inspired by the predictive processing approach to perception [63, 36, 13], yielded in a set of computational models of body illusions that follow the prediction error minimization as the inner mechanism. [35] aimed to investigate how the distance between the rubber and real hands modulates the proprioceptive drift. The work compared the data obtained from a human RHI experiment and a humanoid RHI experiment. It used the learnt generative model of the sensory signals (proprioceptive, visual and visuotactile) to dynamically estimate the location of the real hand. Mathematically, the model used approximated Bayesian inference (i.e., free energy minimization) to infer the body pose given the input signals. Whilst, this work supports the separation of body-ownership illusion and proprioceptive drift, as these seem to be split into two interconnected processes, as suggested by [1, 2], it only provided the computation for replicating the proprioceptive drift and not the body-ownership. In a follow-up model [46, 49], the focus shifted from the passive perceptual process to the active component of the illusion. The model and human evidence suggests that participants actively move their body position (or generate forces) to minimize the discrepancy between proprioceptive signals coming from the real hand, and visual signals coming from the rubber hand. As an aside result, the model explained that perceptual drifts are not necessarily linked to tactile stimulation, but is achievable with visual and proprioceptive signals in VR [50].

The models described above assumed that the location of the hand is already processed. The work by [69] introduced deep neural networks to allow the body illusion model to use large-scale visual inputs. The architecture implemented the deep active inference framework for continuous time variables [72]. This model was able to account for the perceptual and active drifts dynamics during the illusion using synthetic images generated from VR. Furthermore, it introduced the synchronicity of the input signals as a variable. However, this model did not contribute to the biological underpinnings of body ownership. There are other connectivity based models, such as [76], where the neural network connections are inspired by brain functional segregation.

### 5.2. Precision parametrization

Precision learning follows the online learning of measurement noise precision (or the inverse noise covariance matrix) Π^*z*^. Intuitively, this refers to the brain’s attentional mechanism of gauging the right amount of noise in the environment so as to make inferences about the world under high noise (uncertainties)— see [65, 64]. We postulate that an accurate attentional mechanism (online precision learning) is critical to experiencing the RHI. We use the FEP for online precision learning. This follows two steps: modeling and learning. The noise precision Π^*z*^ is modeled through an exponential parametrization with respect to parameter *λ* = [*λ*^1^ *λ*^2^ … *λ*^*m*^]^*T*^ (where *m* is the number of observations) as:

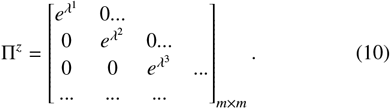

This parametrization was chosen to ensure that Π^*z*^ always remains positive definite (Π^*z*^ ≻ *O*) throughout the estimation. Intuitively, this ensures the noise covariance to be a positive real number.

### 5.3. Code and parameters used for simulations

The MATLAB code ^4^ for the simulation is available: https://github.com/ajitham123/precision_RHI/. This section lists the parameters that were used in our simulations. The generative process uses the model given in Eq (3). The same model is used as *M*_1_ with *Ay* = 0 and *Ry* = 0.8, while the model in Eq (4) is used as *M*_2_. The *α* in Eq (2) is set to 10 for a low sensitivity to the illusion, and to -10 for a high sensitivity to the illusion. The generative process is simulated for 32 seconds with a sampling time of 0.1s. The observation noise for each signal is parametrized using 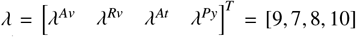 for hand visible, and *λ* = [− 1, 7, 8, 4] for hand covered (see Figure (3) for the generated data). The causes (hand strokes) **X** (plotted in blue in Figure (6)) comprises of i) four Gaussian bumps centered around *t* = 4, 6, 8, 10 seconds for hand visible case and ii) five Gaussian bumps centered around *t* = 20, 22, 24, 26, 28 seconds for hand covered case. The magnitude of these bumps for **Ea, Er, Et, Sy** are 0.5, 1, 1, 0, respectively (when stroked). The estimation starts with priors on noise parameters (in Eq (6)) as *η*^*λ*^ = [10, 10, 10, 10] with prior precision *P*^*λ*^ = *diag*([*e*^−4^, *e*^−4^, *e*^−4^, *e*^−4^]) where *diag*(.) converts an array into a diagonal matrix. All the numbers described here were arbitrarily selected to start the estimation from low noise levels with low confidence. For breaking the precision learning in figure 4, a higher prior precision (*P*^*λ*^ = *diag*([*e*^10^, *e*^−4^, *e*^−4^, *e*^−4^])) for the signal *Av* was used. The state estimation starts at *t* = 0 with a unit precision. All the learning rates in Section 2.2.4 are set to *k* = 4 for a quick learning. All figures are generated using these parameters as the basic reference for the simulation, unless another value is mentioned for a few variables within the text.

## Footnotes

2 In the FEP literature, the generative process refers to the real world process that the agent has only access through the senses, and the generative model refers to the agent model approximation of the reality learnt by experience.

3 See Discussion section for the reason of using model selection instead of model averaging, which is usually followed in computational neuroscience.

4 The code will be made public upon paper acceptance

